# Behavioral context shapes vocal sequences in two anuran species with different repertoire sizes

**DOI:** 10.1101/2021.03.18.436029

**Authors:** Ananda Shikhara Bhat, Varun Aniruddha Sane, Seshadri K S, Anand Krishnan

## Abstract

Acoustic signals in animals serve to convey context-dependent information to receivers. Birds and mammals combine diverse sounds into complex sequences to communicate, but these sequences largely remain understudied in other taxa. Anuran vocalizations are a prominent feature of their life history, and function in defense of territories and to attract mates. However, despite the spectacular diversity of anurans in tropical regions of the world, vocal diversity and communication strategies remain relatively poorly studied. Specifically, studies of vocal sequences and context-dependent vocal patterns in frogs remain few. Here, we investigated the context-dependent vocal repertoire and the use of vocal sequences by two anuran species belonging to different lineages, both endemic to the hyper-diverse Western Ghats of India. By recording vocal sequences both when frogs were alone and in the presence of a territorial rival, we present evidence that both species modify their vocal repertoire according to context. Specifically, one species appends notes to generate more complex sequences, whereas the other shifts to different note types, resulting in different sequences for different contexts. Thus, despite differences in repertoire size, both frog species are capable of adjusting the temporal sequence of vocalizations to communicate in different contexts. This study highlights the need for further studies of insular frogs, to understand how diversification across these continental islands has influenced the evolution of vocal repertoires, vocal sequence patterns and communication systems.

**Lay Summary:** Animals employ complex sequences of acoustic signals to communicate in diverse behavioral contexts. Here, we demonstrate that two frog species with different vocal repertoires both modify the sequence of note emissions in the presence of a territorial rival. These patterns demonstrate that anurans are capable of complex shifts in the patterns of their vocalization, to communicate different messages to different receivers. Our findings demonstrate the value of studying behavioral diversity in tropical regions.

## Introduction

Acoustic communication has evolved in a variety of taxa (Bradbury and Vehrencamp 2011; Chen and Wiens 2020), and enables different messages to be conveyed to different receivers, often simultaneously. The production of sound is energetically costly (Gerhardt 1994; Gillooly and Ophir 2010) and also carries the risk of detection by unintended receivers such as predators or parasites (Wells 1977a; Rand and Ryan 1981; Zuk et al. 2006). These factors compound the challenge of conveying information effectively to intended recipients, such as prospective mates, or territorial rivals, among others (Wilkins et al. 2013). To solve this problem, animals may evolve signals or signaling behavior to suit each behavioral context, for example, employing distinct acoustic signals for a mate versus a rival (Berglund et al. 1996; Hebets and Papaj 2005). These may result in complex temporal patterns in acoustic signals, and in some animals, receivers may be able to distinguish these sequences from each other (Gentner et al. 2006; Fishbein et al. 2020). However, study of animal communication using complex vocal sequences remains limited, and most studies have focused on birds and mammals (Marler 1990; Gentner et al. 2006; Bohn et al. 2009; Kershenbaum et al. 2012; Allen et al. 2018; Fishbein et al. 2020). Thus, temporal patterning and the use of sequences in the vocalizations of other taxa remain poorly understood.

Anurans (frogs and toads) are a diverse amphibian order that have evolved conspicuous acoustic signals to attract females (Narins 1992; Narins 1995), which form one of the most prominent features of their life histories. In addition to serving as honest indicators of mate quality, anuran acoustic signals are also used to advertise position, convey information about reproductive state, defend territories, and communicate fitness to rivals of the same sex (Wells 1977b; Bradbury and Vehrencamp 2011; Bee, Schwartz, et al. 2013). Although studies have sought to ascribe call types to different functions (Toledo et al. 2015), there is still little knowledge of whether anurans employ vocal sequences of different note types, and whether the sequence of vocalizations varies according to behavioral context. In tropical regions, which possess hyper-diverse amphibian communities (Bossuyt et al. 2004; Jetz and Pyron 2018), there is a severe paucity of basic information on frog vocal repertoires, and virtually no studies on vocal sequences or vocal behavior (Aravind and Gururaja 2011). The diversity of signals and signaling strategies that potentially exist in these areas may help identify broader principles underlying context-dependent acoustic signaling and complex communication in amphibians. Comparative studies of tropical amphibians are also particularly relevant in light of global amphibian declines (Houlahan et al. 2000; O’Hanlon et al. 2018), and the discovery of new species and behaviors from the tropics (Diesel et al. 1995; Biju and Bossuyt 2003; Feng et al. 2006; Shen et al. 2008; Gururaja et al. 2014; Iskandar et al. 2014; Seshadri et al. 2015; Vijayakumar et al. 2019).

Here, we examine context-dependent vocal behavior in two anuran species, representatives of endemic lineages from India’s hyper-diverse Western Ghats. *Nyctibatrachus* and *Pseudophilautus* have both evolved in the Western Ghats-Sri Lanka biodiversity hotspot (Van Bocxlaer et al. 2012; Vijayakumar et al. 2016; Meegaskumbura et al. 2019; Torsekar 2019). Although studies of anuran vocalizations from this region have characterized taxonomic diversity (Garg et al. 2021), relatively little remains known about vocal behavior. Members of both lineages use vocalizations to communicate and are prolonged, territorial breeders (Gramapurohit et al. 2011; Bee, Suyesh, et al. 2013). They are thus likely to communicate both with potential mates and with territorial rivals. Our study a) quantifies the diversity of their vocal repertoires, and b) examines the use of notes and note sequences in different behavioral contexts. We find that these two species exhibit different repertoire sizes, and proceed to address the question of how males adapt these repertoires to territorial contexts, specifically the presence of a rival male.

## Materials and Methods

### Study Species

We studied two endemic anuran species, *Nyctibatrachus humayuni* (Nyctibatrachidae) and *Pseudophilautus amboli* (Rhacophoridae) at distinct locations in India’s Western Ghats. Both species are relatively common within their geographic ranges (Van Bocxlaer et al. 2012; Vijayakumar et al. 2016; Meegaskumbura et al. 2019). Humayun’s Night frog (*Nyctibatrachus humayuni*) (Bhaduri and Kripalani 1954) is a relatively large frog restricted to the Northern Western Ghats (males measuring up to 46.8 mm and females up to 50.6 mm), found in forest streams and swamps between 560-1228 metres above sea level (Fig 1). Males of this species are territorial, and vocalize at night on rocks or on vegetation overhanging water. The Amboli bush frog, (*Pseudophilautus amboli*) (Biju and Bossuyt 2009) is a medium-sized frog restricted to the Northern and Central Western Ghats (males measuring up to 34.1 mm and females up to 37.5 mm). Individuals vocalize at night on leaves and saplings at heights of up to two meters from the ground in a variety of habitats, including disturbed evergreen forests, arecanut (*Areca catechu*) plantations, and gardens between 500-1000 metres above sea level (Fig 1). Both species are easy to locate in their preferred habitats, and are thus relatively easily observed and recorded. They thus present an opportunity to relate vocalizations and vocal sequence patterns to behavioral context.

**Figure 1:**
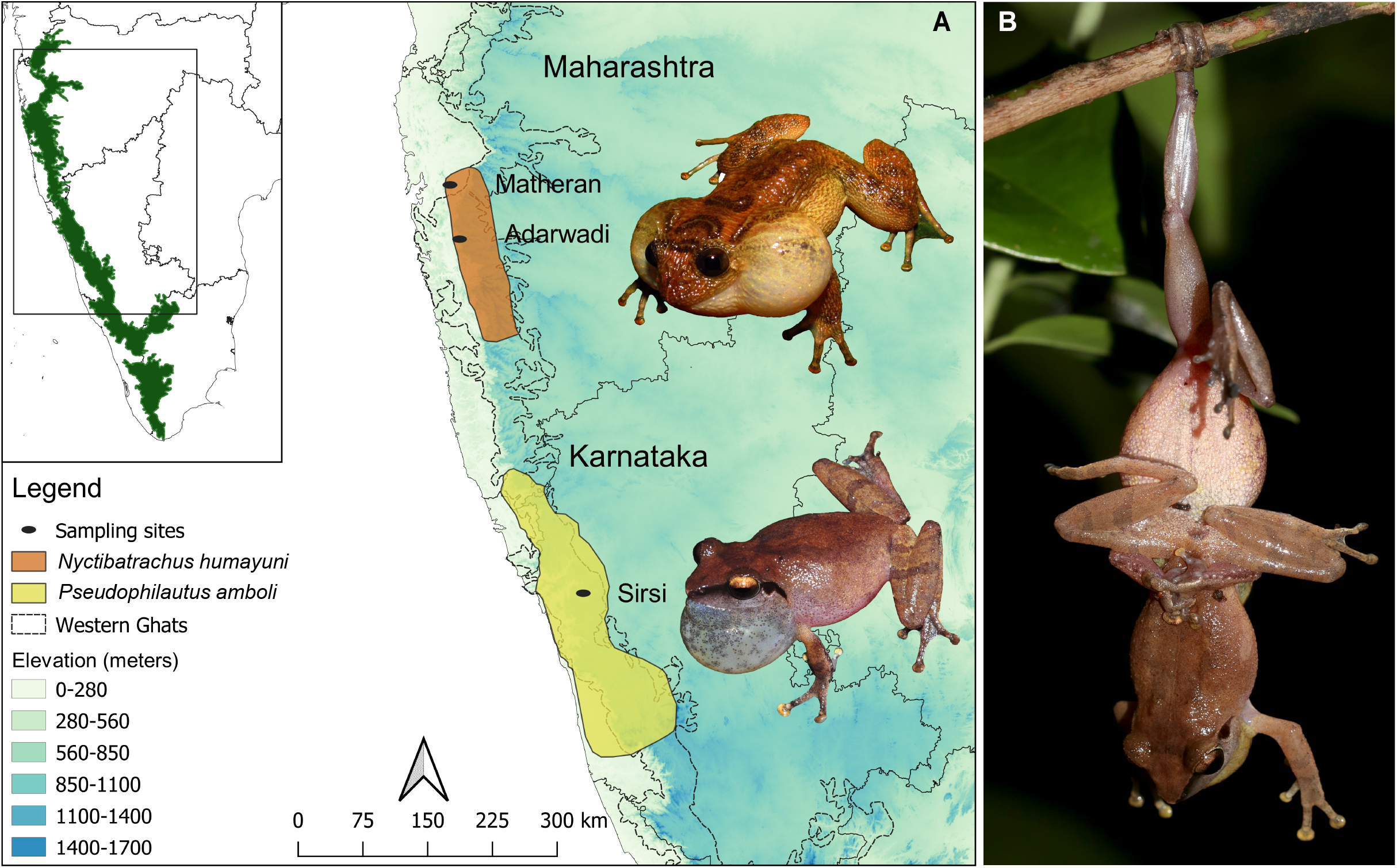
**A.** Geographical distributions of *Nyctibatrachus humayuni* (top) and *Pseudophilautus amboli* (bottom). Inset map shows the Western Ghats in green and the box indicates the study area. **B.** Two male *P. amboli* engaged in physical male-male combat.

### Study sites

We studied *N. humayuni* at two locations in Maharashtra (Adarwadi village, Pune district, 18° 25.838’N, 73° 24.085’ E, 634 metres above sea level, and Matheran, Raigad district, 19°00’N, 73°17’E, 800 metres above sea level) in June-July 2019. Matheran is a known locality for *N. humayuni* (Biju et al. 2011; Gramapurohit et al. 2011; Joshi et al. 2019). These localities comprise mid-elevation wet evergreen forest, receiving about 4073 mm rainfall annually (Sawant et al. 2020). We studied *P. amboli* in Sirsi, Uttara Kannada district, Karnataka (14°44’N, 74°45’E, 611 metres above sea level) in July 2020. This locality comprises a semi-evergreen forest (http://wgbis.ces.iisc.ernet.in/energy/water/paper/ETR24/index.htm) and receives a mean annual rainfall ranging between 2500-3500 mm with peak rainfall during the months of June-August (Bhat 1992).

### Recording vocalizations

At each site, we recorded frogs in clear weather between 18:30 and 22:00 hours, as both species are nocturnal or crepuscular. For recording, we used a Sennheiser (Wedemark, Germany) microphone connected to a Zoom (Tokyo, Japan) H6 recorder (*N. humayuni*) or a Marantz (Kawasaki, Kanagawa, Japan) PMD-660 recorder (*P. amboli*). We recorded at least 20-30 calls per frog at a sampling rate of 44.1 kHz. Before beginning a recording session, we assigned individuals to behavioral context as follows: **Alone** (a single vocal frog with no vocalizing neighbors within an estimated 100 body lengths, i.e. snout-vent lengths in the horizontal plane); **Vocalizing with neighbors** (another vocal male within 100 body-lengths in the horizontal plane: we only considered cases where the vocal neighbor was visible to us, so as to be stringent in our criteria); and **Territorial dispute** (a confrontation between vocal males, including aggressive posturing and physical contact). We used 100 body-lengths in the horizontal plane as a measure based on published studies of nearest neighbor distances (Kunte 2004). All social contexts involve only a single neighbor, as incidents with more than one vocal neighbor within this radius were extremely rare (and have therefore not been analyzed as part of this dataset). We only analyzed vocal bouts where we could keep track of the frog’s movements throughout the period of observation, and took care to sample different localities or territories on different nights to avoid recording the same male twice.

### Analysis

Using Raven Pro 1.6 (Cornell Laboratory of Ornithology, Ithaca, NY, USA), we digitized individual call notes, and classified them into various note types based on their time-frequency properties. For further analysis, we used custom codes written in Python and R (R Core Team 2020), as well as the Python packages numpy (Harris et al. 2020), seaborn (Waskom 2020), matplotlib (Hunter 2007), pandas (McKinney 2011) and scipy (Virtanen et al. 2020), and the R packages ggplot2 (Wickham 2016), ggpubr (Kassambara 2020), stringr (Wickham 2019) and dplyr (Wickham et al. 2021).

First, we calculated the number of each note type emitted by each individual, as a proportion of the total notes (of all types). We compared these proportions across contexts for each note type using either a Mann-Whitney U-test (*N. humayuni*, where we compared two contexts), or a Kruskal-Wallis test (*P. amboli*, where there were three contexts). Our null hypothesis in either case was that the proportion of a particular note type was similar across contexts, i.e. that there were no context-dependent changes in the use of specific note types.

For *P. amboli*, we performed additional analyses to further examine context-dependent changes in the sequences of vocalization. First, we examined note diversity or variability in the vocal repertoire during each context using Shannon entropy. The Shannon entropy of a stochastic process with n different possible states is given by 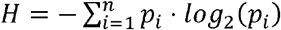, where p_i_ is the probability of occurrence of the i^th^ state, and if p_i_ = 0, we define log_2_(p_i_) = 0. H is a positive number between 0 and log_2_(n), and a higher value of H indicates that the process is more ‘unpredictable’. In information-theoretic terms, a higher value of H corresponds to a process that carries more information (Shannon 1948; Kershenbaum et al. 2016). In other words, a stochastic process with a higher value of H has a higher ***diversity*** of occurrence of states. For frog vocalizations, therefore, a **higher** value of H corresponds to **greater note diversity**, or more variation in vocal sequences in any given behavioral context. Using custom scripts, we calculated the Shannon entropy for each individual for each context, and compared across contexts using a Kruskal-Wallis test. Next, we examined changes in vocal sequences across contexts. To accomplish this, we modelled vocalizations as a first-order Markov chain to calculate the transition probabilities between different note types, where each vocalization depended only on the one immediately preceding it. For a time-homogeneous, discrete-time, discrete-space Markov chain with n distinct states (in our case, the number of different note types) and the random variable X_t_ (corresponding to the value of note number t, or the note under consideration), the process provides an n x n transition probability matrix *P* = [*p_ij_*], (where *p_ij_* = *Pr{X_t+1_* = *j |X_t_ = i}*). Using maximum likelihood estimation and Lagrange multipliers for a first-order Markov chain, p_ij_ can be estimated from data as:

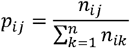

Where *n_ij_* denotes the number of times an *i → j* transition occurred.

This gives us the probability that each note type is followed by another note type, and we examined these to understand how the sequences of different note types changed across contexts.

We calculated transition probabilities for each context using the above-mentioned custom scripts. A string of the sequence of calls emitted by an individual was first extracted from the data. In this way, we created strings for all individuals for each context. The transition matrix P was then calculated by estimating each n_ij_ for each context. We additionally cross-verified the results from the Markov chain analysis by performing a second analysis of note sequences, because animal vocal sequences may not always follow Markovian dynamics (Kershenbaum et al. 2014). This second analysis made no assumptions about the underlying stochastic process used to generate the observed vocal sequences. For a given set of note sequences, we defined ^d^C_ij_ as the probability that notes ‘i’ and ‘j’ occur within a distance of ‘d’ notes from each other. Using a Monte-Carlo sampling method, we first calculated ^d^C_ij_ for the vocal sequence data from our recordings. We then constructed sequences for comparison, where notes were ‘randomly’ distributed according to a stationary uniform distribution (i.e. probability of emitting a given note-type was equal to the proportion of that note-type in the dataset). In these latter sequences, the probability of occurrence of each note was equal to its proportion of occurrence (# of focal note type/total # of all note types combined) in the dataset. We synthesized 50 such random datasets and averaged the co-occurrence across all of them to obtain a robust estimate of the expected (“null”) co-occurrence. Comparing the ratio ^d^R_ij_ of the observed to expected co-occurrence allowed us to quantify whether the notes ‘i’ and ‘j’ occur within a distance of ‘d’ notes more or less often than we would expect in a random sequence of notes. A value of ^d^R_ij_ < 1 means that the notes ‘i’ and ‘j’ occur within a distance of ‘d’ less often than random, and ^d^R_ij_ > 1 means that the notes ‘i’ and ‘j’ occur within a distance of ‘d’ more often than random. We carried out this analysis using different values of d, to assess how the patterns depended on the length of sequence under consideration (See Supplementary figure 01 for an illustration of the workflow).

## Results

### Vocal repertoires in endemic anuran lineages

Both species of anurans emitted calls as sequences comprising two or more note-types. Specifically, *N. humayuni* produced vocal sequences comprising two note types (Figure 2A). According to existing classification systems (Emmrich et al. 2020), *N. humayuni* falls within call guild F (“frequency modulated call with uniform notes”). Henceforth, the first note type is referred to as an ‘Ascending Note’ (AN; note emission rate = 5.425 ± 2.360 notes/minute; note duration = 370 ± 66 ms, range = 228 - 619 ms; dominant frequency = 1575.2 ± 237.58 Hz, range = 1033.6 - 2153.3 Hz; N = 489 notes from 19 males). The AN was the more commonly recorded of the two, constituting 66.2% of total notes recorded across individuals. The second note type, which we refer to as the ‘Descending Note’ (DN), was lower in frequency (note emission rate = 2.681 ± 2.450 notes/minute; note duration = 360 ± 57 ms, range = 230 - 553 ms; dominant frequency = 1098.0 ± 304.86 Hz, range = 602.9 - 1722.7 Hz; N = 243 notes from 16 males). DNs were typically emitted in variable numbers immediately following an AN. Thus, a typical call followed the sequence (AN + x*DN), where x ranged between 0 and 8.

**Figure 2:**
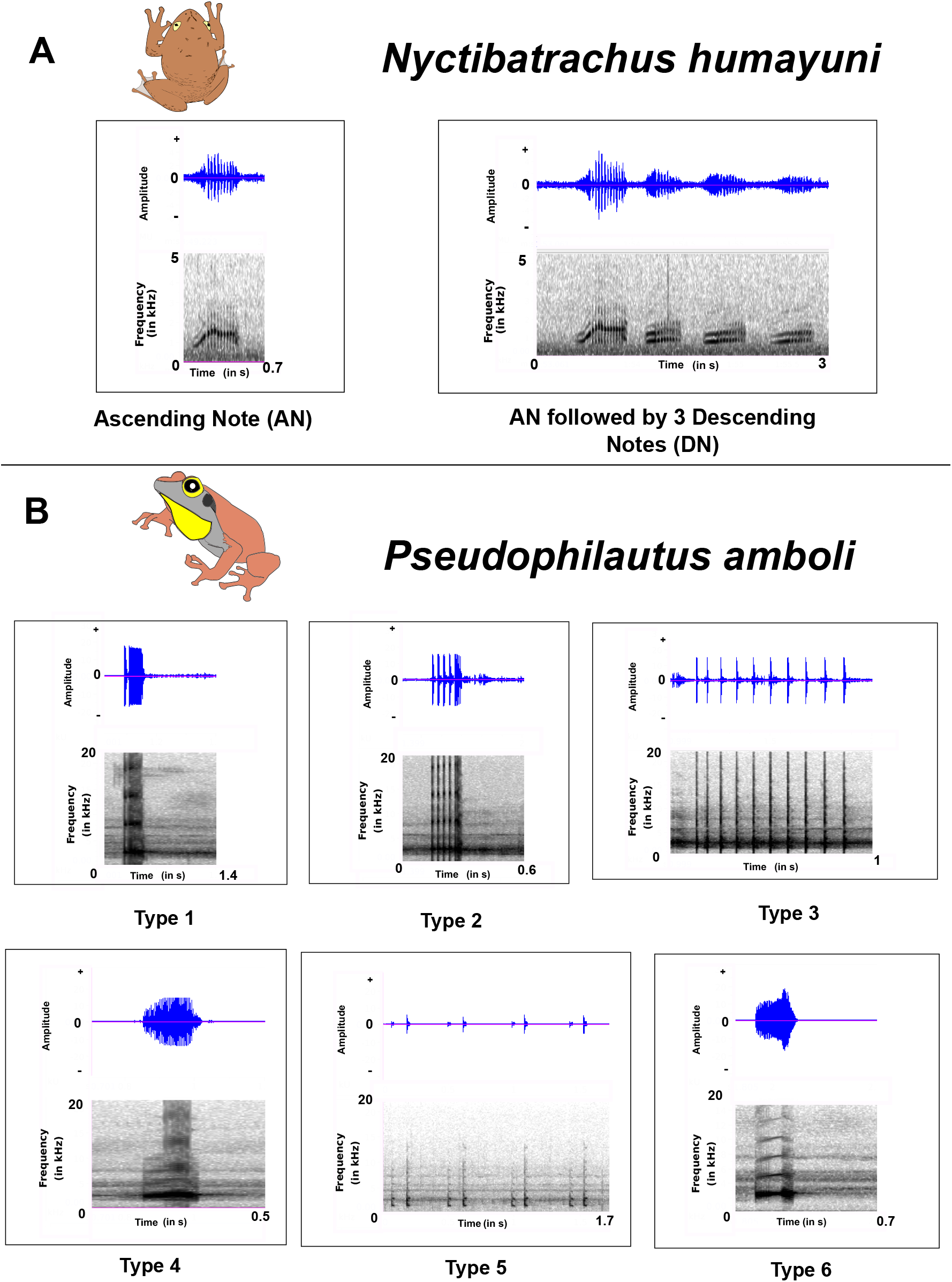
Oscillograms and spectrograms of the vocal repertoire of **A.** *N. humayuni* and **B.** *P. amboli*

Males of *P. amboli* fell within call guild G (“non-frequency-modulated complex calls”). The vocal repertoire of this species consisted of 6 distinct note types (henceforth referred to as note types 1-6; note parameters in Table 1; spectrograms in Figure 2B). Among all calls recorded, note type 5 was most frequently encountered, comprising 28.7% of total recorded notes, whereas the rarest was note-type 6, comprising only 1.02% (this note type was recorded too infrequently for statistical analysis, and thus was not considered when calculating transition probabilities). Individuals emitted between 2 to 6 different note types in a single recording session.

**Table 1:** Properties of the note-types of *P. amboli*

| Note-type | Peak Frequency<br>(Hz) and SD | Note duration<br>(ms) and SD | Bandwidth 90%<br>(Hz) |
| --- | --- | --- | --- |
| 1 | 2669.9 ± 131.28 | 154.4 ± 40.1 | 2859.3 |
| 2 | 2610.7 ± 162.32 | 218.8 ± 44.6 | 2857.7 |
| 3 | 2283.7 ± 536.02 | 48.7 ± 11.7 | 2544.1 |
| 4 | 2478.0 ± 382.11 | 312.6 ± 105.4 | 1106.7 |
| 5 | 1954.4 ± 483.61 | 49.4 ± 16.1 | 1532.0 |
| 6 | 2557.5 ± 62.96 | 219.5 ± 55.1 | 439.5 |

### N. humayuni emphasizes DNs in territorial contexts

Vocalizing males of *N. humayuni* varied the proportion of DNs in vocal sequences, appending more DNs following an AN in the presence of a vocalizing neighbor (Wilcoxon rank-sum test: *W* = 8, N = 19, p < 0.01. Figure 3A). However, they did not vary the note emission rate (Wilcoxon rank-sum test, *W* = 49, N = 19, p > 0.5). When vocalizing alone, males often emitted only the AN, and only rarely added DNs following it. In most cases where a neighbor was present, frogs typically appended a single DN to their call (Figure 3B), and rarely up to 3 DNs, thus resulting in a more complex vocal sequence. We observed only a single ‘territorial dispute’ involving physical contact between two males. In this encounter, the male under observation regularly emitted high numbers of DNs, ranging up to 8 DNs following an AN (Supplementary Audio 01).

**Figure 3:**
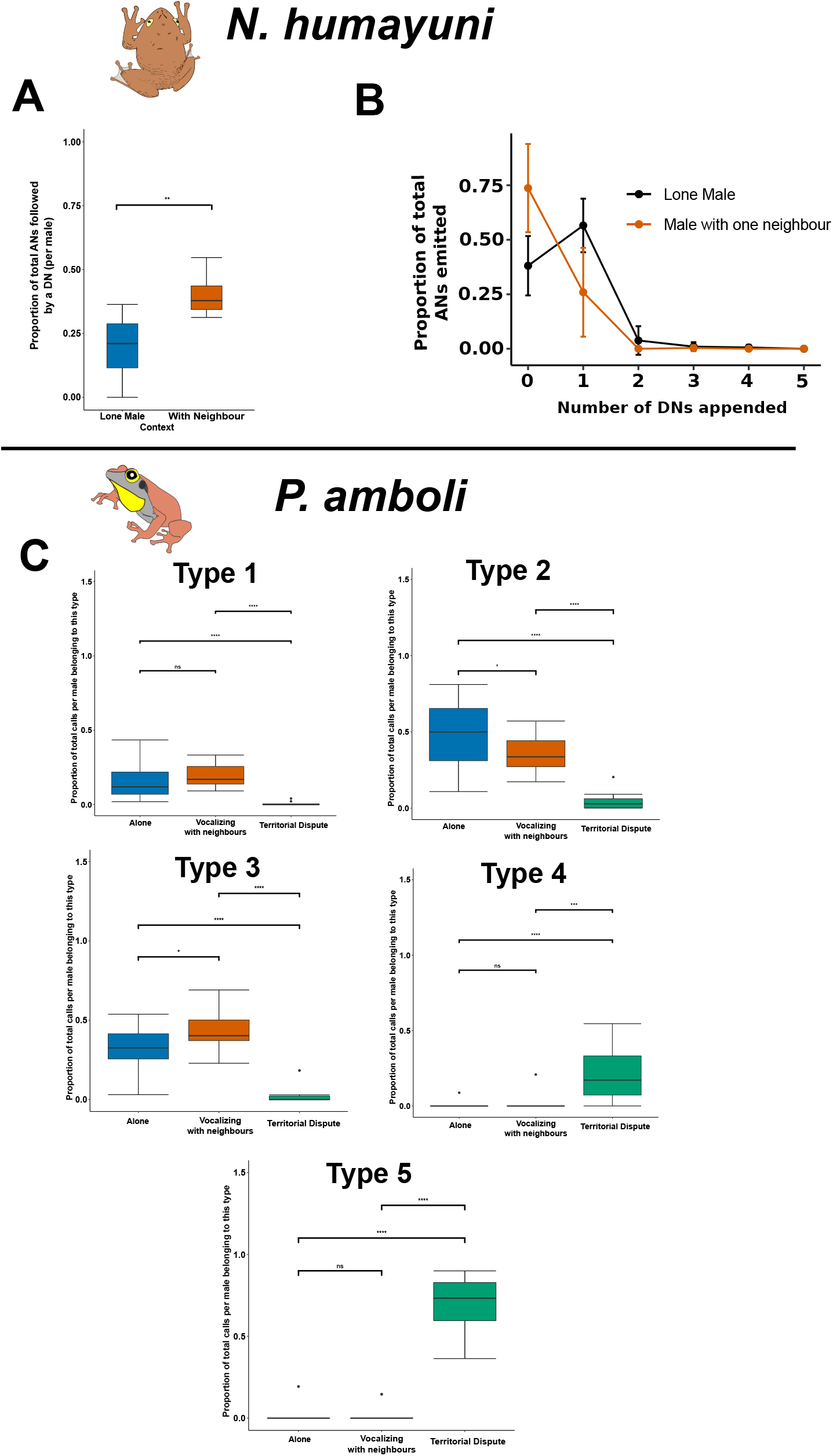
**A.** Variation of proportion of each note type used across behavioral context in *N. humayuni*. **B.** Proportion of times a male *N. humayuni* appended different numbers of DNs in each context. **C.** Variation of proportion of each note type used across behavioral contexts in *P. amboli* (****-p<0.0001, ***-p<0.001, **- p<0.01, *- p<0.05, ns- not significant).

### P. amboli utilizes different vocal sequences in different behavioral contexts

Male *P. amboli* varied the proportions of each note type in vocal sequences, according to behavioral context (Figure 3C, Table 2). Males vocalizing either alone or with a neighbor emitted note types 1, 2 and 3 most often, whereas individuals engaged in territorial disputes mostly emitted note types 4 and 5. Furthermore, males vocalizing with neighbors emitted a higher proportion of type 3 notes (Kruskal-Wallis test: *H_1_* = 4.72, p < 0.05) and a lower proportion of type 2 notes (Kruskal-Wallis test: *H_1_* = 5.02, p < 0.05) than males vocalizing alone.

**Table 2:** Variation of proportion of note-types of *P. amboli* based on behavioral context (Results of Kruskal-Wallis test)

| Note-Type | Results of Kruskal-Wallis test |
| --- | --- |
| 1 | $H_2 = 24.55, p = 4.7 \times 10^{-6}$ |
| 2 | $H_2 = 26.04, p = 2.2 \times 10^{-6}$ |
| 3 | $H_2 = 26.67, p = 1.6 \times 10^{-6}$ |
| 4 | $H_2 = 32.03, p = 1.1 \times 10^{-7}$ |
| 5 | $H_2 = 41.11, p = 1.2 \times 10^{-9}$ |

These differences in use of note types were accompanied by differences in Shannon entropy based on social context (Kruskal-Wallis test: *H*_2_ = 25.38, p < 0.001. Figure 4A). Males engaging in territorial disputes exhibited a lower Shannon entropy, which is indicative of more stereotyped vocal sequences. When we modelled vocal sequences in each context as a first-order Markov chain, we observed differences in the transition probabilities of different note types based on context (Figure 4B). These transition probabilities resulted in grouping within note types, regardless of the initial state of the system. The groupings of note types followed the same context-dependent patterns described above (circle sizes in Figure 4B).

**Figure 4:**
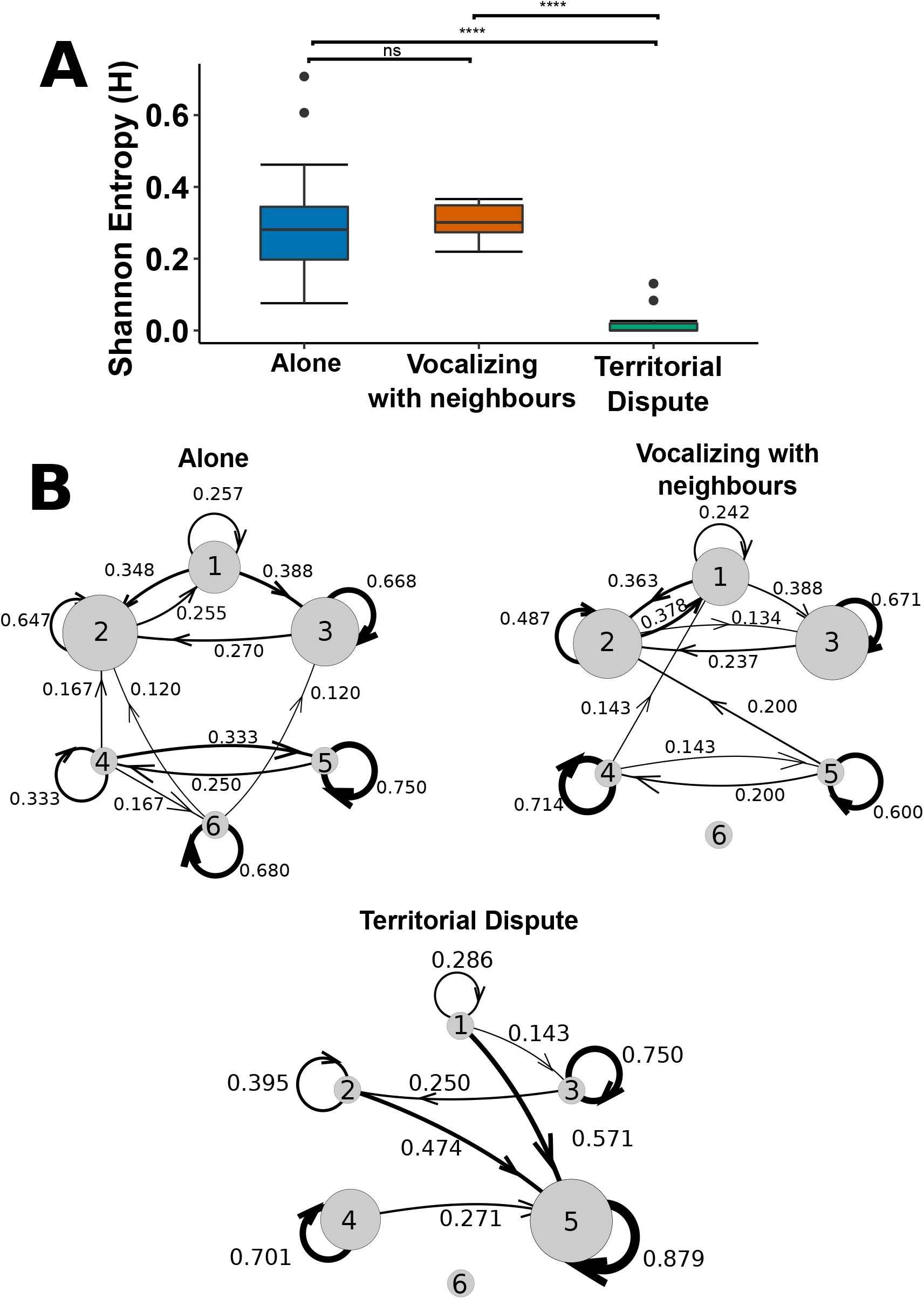
**A.** Variation in Shannon entropy of vocalization with behavioral context in *P. amboli* (****- p < 0.0001, ***-p < 0.001, **- p < 0.01, *- p < 0.05, ns- not significant). **B.** Transition probabilities obtained by modelling the vocal sequence of *P. amboli* as a first-order Markov chain for different behavioral contexts. Size of each circle represents the average proportion of that note type (relative to all notes produced) in that context, and numbers on the arrows are transition probabilities. Transition probabilities that are less than 0.1 have been omitted for the sake of clarity.

Using Monte-Carlo methods to calculate the probability of two note types occurring within d = 6 notes of each other (the size of the repertoire for this species), we were able to verify the conclusions from the Markov chain analysis. Plotting the pairwise measures of the ratio of observed to expected probability of co-occurrence (see Methods) uncovered two note groups, one containing note types 1, 2, and 3, and the second containing note types 4 and 5. (Figure 5). ^d^R_ij_ was > 1 for intragroup pairings, and < 1 for intergroup pairings. This trend was independent of the value of d (3 <= d <= 12, additional plots in Supplementary file 01). The presence of note groups, together with the results of the Markov analysis, are consistent with the presence of different vocal sequences for different contexts. Male *P. amboli* thus shift from one note group to the other in a territorial dispute, resulting in more stereotyped vocal sequences.

**Figure 5:**
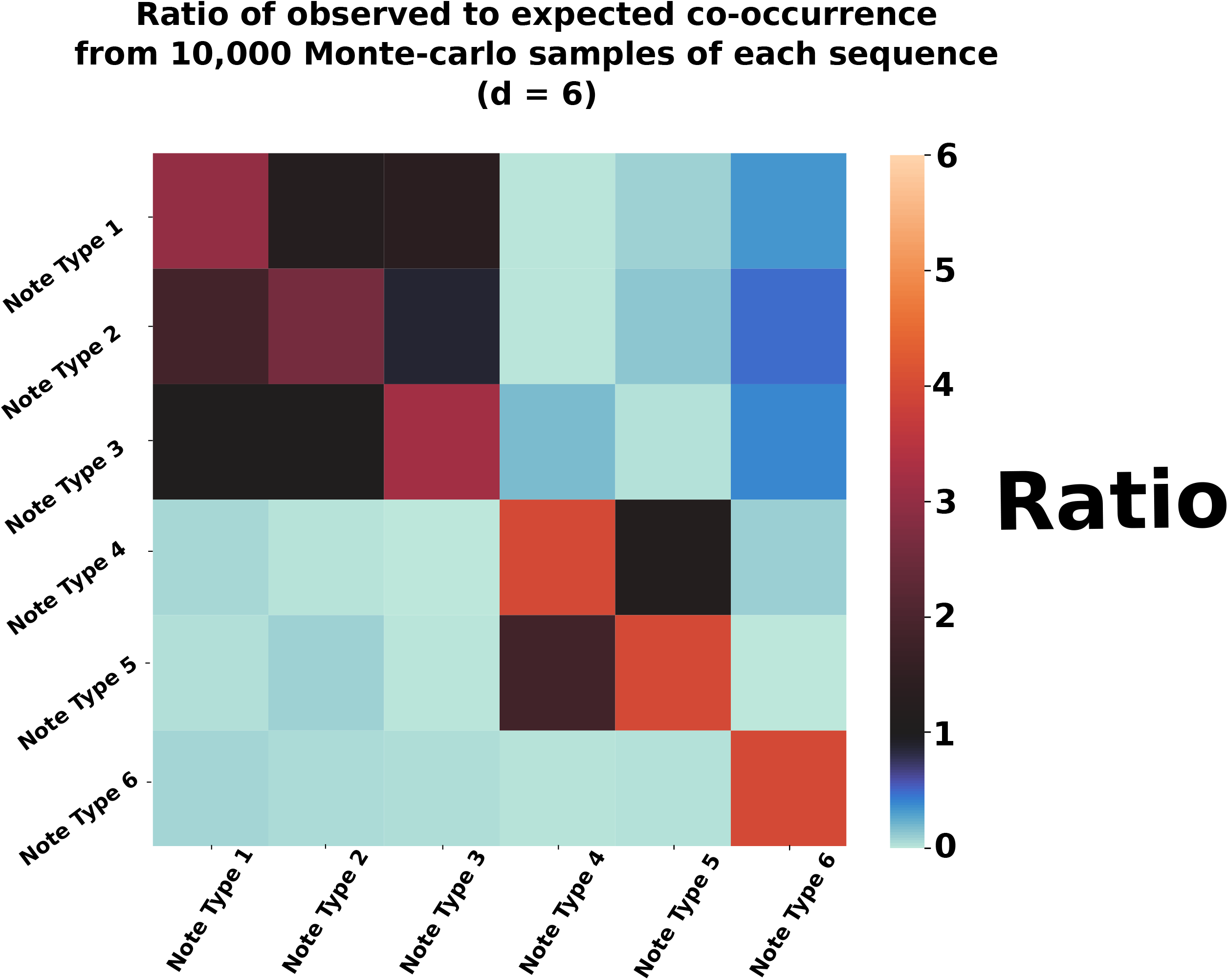
Ratio of observed to expected probabilities of note co-occurrence (^6^R_ij_), for *P. amboli* calculated using 10,000 runs of the Monte Carlo method for each sequence, with expected co-occurrence being averaged over fifty synthesized random datasets. Color indicates the probability that two notes co-occur in a sequence within 6 notes of each other. Redder colors indicate that the corresponding note types co-occur more often than expected by chance alone.

## Discussion

In summary, we uncover evidence that members of two distantly related frog lineages, endemic to India’s Western Ghats, solve the challenge of context-dependent communication in broadly similar ways. Both species (with different repertoire sizes) modify their vocal sequences in the presence of a territorial rival by incorporating different note types. One species appends notes, resulting in more elaborate sequences, and the other switches to a different group of notes in the presence of a rival, resulting in more stereotyped sequences of notes. Thus, the different note types in anuran repertoires may be emitted in sequences that support context-dependent communication.

Signalers of many taxa communicate both with potential mates and territorial rivals (Wells 1977b; Johnstone 1996; Candolin 2003; Hebets and Papaj 2005). In some cases, the signaler may employ entirely different signals for different receivers, or in different contexts (Leo 1959; Baptista 1978; Balakrishnan and Pollack 1996; Andersson et al. 2002; Byers and Kroodsma 2009; Karubian et al. 2009; Vanderbilt et al. 2015; Zambre and Thaker 2017; Rosenthal et al. 2018; Centeno et al. 2020). Several frog species across the world possess ‘compound advertisement calls’ in which one component is directed at other males, and the other is directed at females (Wells and Greer 1981; Arak 1983; Schwartz and Wells 1984; Littlejohn and Harrison 1985; Backwell 1988; Nali and Prado 2014; Furtado et al. 2016). The two components of the call are thus ‘functionally partitioned’. For example, *Eleutherodactylus coqui* possesses a two component ‘co-qui’ call, where the ‘co’ component is directed towards males, and the ‘qui’ component is directed towards females (Narins and Capranica 1978).

Alternatively, signalers with limited repertoires may modify the sequence in which notes are emitted, such that similar signals may convey different messages to different receivers (Berglund et al. 1996; Myberg Jr 1997; Dalziell and Cockburn 2008; Vasconcelos et al. 2010; Moskát and Hauber 2019). A notable example of such a strategy is seen in the túngara frog (*Engystomops pustulosus*) wherein the male vocal repertoire comprises two notes: a ‘whine’ and a ‘chuck’. A male may emit only a whine (a ‘simple’ call), or append a variable number of chucks to the end of a whine to produce a ‘complex’ call (Rand and Ryan 1981). Females prefer complex calls over simple calls (Ryan 2019); however, vocal males adjust the complexity of their own call in response to other complex calls (Ryan 2019). Therefore, multi-note vocal repertoires in anurans (Narins et al. 2000) present a number of avenues by which vocal sequence may be modified according to behavioral context.

We demonstrate that the presence of conspecific males drives changes in vocal sequence in both frog species. In *P. amboli*, note types 4 and 5 are emitted only during territorial disputes, resulting in greater sequence stereotypy. This suggests an ‘agonistic’ or ‘territorial’ function for sequences consisting of these notes, directed towards other males. Note type 1, on the other hand, was emitted only in non-dispute contexts, and did not vary with the presence of conspecific males in the area. We thus hypothesize that note type 1 may serve as an advertisement call. Thus, *P. amboli* appears to actively switch to different groups of notes, resulting in very different vocal sequences in different contexts (Figure 4, Figure 5). On the other hand, *N. humayuni* emits the same call types when alone as well as in the presence of other males, owing to a relatively limited repertoire of just two note types. However, in the presence of a neighbor, males append more DNs to the end of their call, resulting in more elaborate vocal sequences. Thus, using the same signals, males of this species modify the sequence of notes to signal to both potential mates and territorial rivals.

Acoustic signals are structured by evolutionary history, physiology and the effects of body size (Ryan 1988). The two anuran lineages containing *Nyctibatrachus* and *Pseudophilautus* radiated independently in the Western Ghats (Van Bocxlaer et al. 2012; Vijayakumar et al. 2016; Meegaskumbura et al. 2019; Torsekar 2019); however, the vocal repertoire of *P. amboli* is larger than that of *N. humayuni*. Furthermore, the two notes present in *N. humayuni* are sequenced in a predictable, stereotyped manner (AN + x*DNs), whereas the 6 note types present in *P. amboli* are strung together in complex, relatively unpredictable vocal sequences. Notwithstanding these differences, changing the sequence of notes enables both species to communicate effectively to their intended recipient.

Acoustic signals play a vital role in mate attraction, species recognition and defense of territories (Narins 1995; Bradbury and Vehrencamp 2011; Wilkins et al. 2013). Communication signals may be predicted to exhibit diversity in support of these varied functions. In hyper-diverse regions such as the Western Ghats, understanding and quantifying this behavioral diversity may provide both scientific insight and important inputs for conservation (Garg et al. 2021). Recent literature suggests that behavioral diversity must be considered when assessing biodiversity for conservation (Caro and Sherman 2012; Cooke et al. 2014; Cordero-Rivera 2017). Anurans in India remain underrepresented in behavioral studies, and their vocal repertoires remain largely undocumented despite being threatened by a range of factors from habitat loss to chytridiomycosis (Dahanukar et al. 2013; Mutnale et al. 2018). We demonstrate that amphibians in this region and other similar hyper-diverse regions may provide useful comparative insight into the function of vocal sequences and the evolution of context-dependent communication. By employing well-defined metrics to characterize vocal sequences quantitatively across behavioral contexts, our study demonstrates how these sequences may be used to communicate with different receivers. This serves as a starting point to further quantify anuran behavioral diversity across the Western Ghats-Sri Lanka biodiversity hotspot. Comparative analysis of vocal behavior across amphibian lineages, including using playback to study female perception and mate choice, could be insightful in understanding the evolution of communicative complexity and diversity in the tropics.

## Acknowledgements

AK and SKS are funded by INSPIRE Faculty Fellowships from the Department of Science and Technology, Government of India, and AK is also funded by an Early Career Research Grant (ECR/2017/001527) from the Science and Engineering Research Board (SERB), Government of India. ASB was additionally funded by a Kishore Vaigyanik Protsahan Yojana (KVPY) Fellowship from the Government of India. Ramdas Yenpure, Amatya Sharma, and Himanshi Kaushik provided logistical support, Akshay Bharadwaj and Sarthak Malusare helped during fieldwork in Sirsi and Matheran respectively. Varad Giri provided us with locations to record *N. humayuni* in Matheran and Gururaja KV provided a photograph of *N.humayuni* for Figure 1. Finally, Shivam Chitnis, and Vidisha Kulkarni provided helpful comments and suggestions.

## Data Availability

All data and scripts used in the analyses reported in this article will be uploaded to a GitHub repository prior to publication.

---

**Supplementary figure 01:** Workflow used to obtain the co-occurrence matrices. In the algorithm on the right, p_i_ denotes the number of times the i^th^ note type was found in a sub-sequence, and N_ij_ denotes the number of times the i^th^ and j^th^ note types were found in the same sub-sequence. For the expected co-occurrence, the average co-occurrence obtained over 50 synthesized (‘random’) datasets was used.

**Supplementary file 01:** R_ij_ matrices for d values of 3, 6, 9 and 12, from 10,000 Monte Carlo samples of each sequence. Redder colors indicate that the corresponding note types co-occur more often than expected by chance alone.

**Supplementary video 01:** Vocal repertoires of *P. amboli* and *N. humayuni* (with spectrograms and oscillograms). Multiple notes of note types 3 and 5 of *P. amboli* are shown, as the duration of a single note is very small.

**Supplementary audio 01:** Male *N. humayuni* emitting an AN followed by 8 DNs during a territorial dispute.

