## Supplementary figures and images for "Behavioral context shapes vocal sequences in two anuran species with different repertoire sizes"

### Supplementary Figure 01

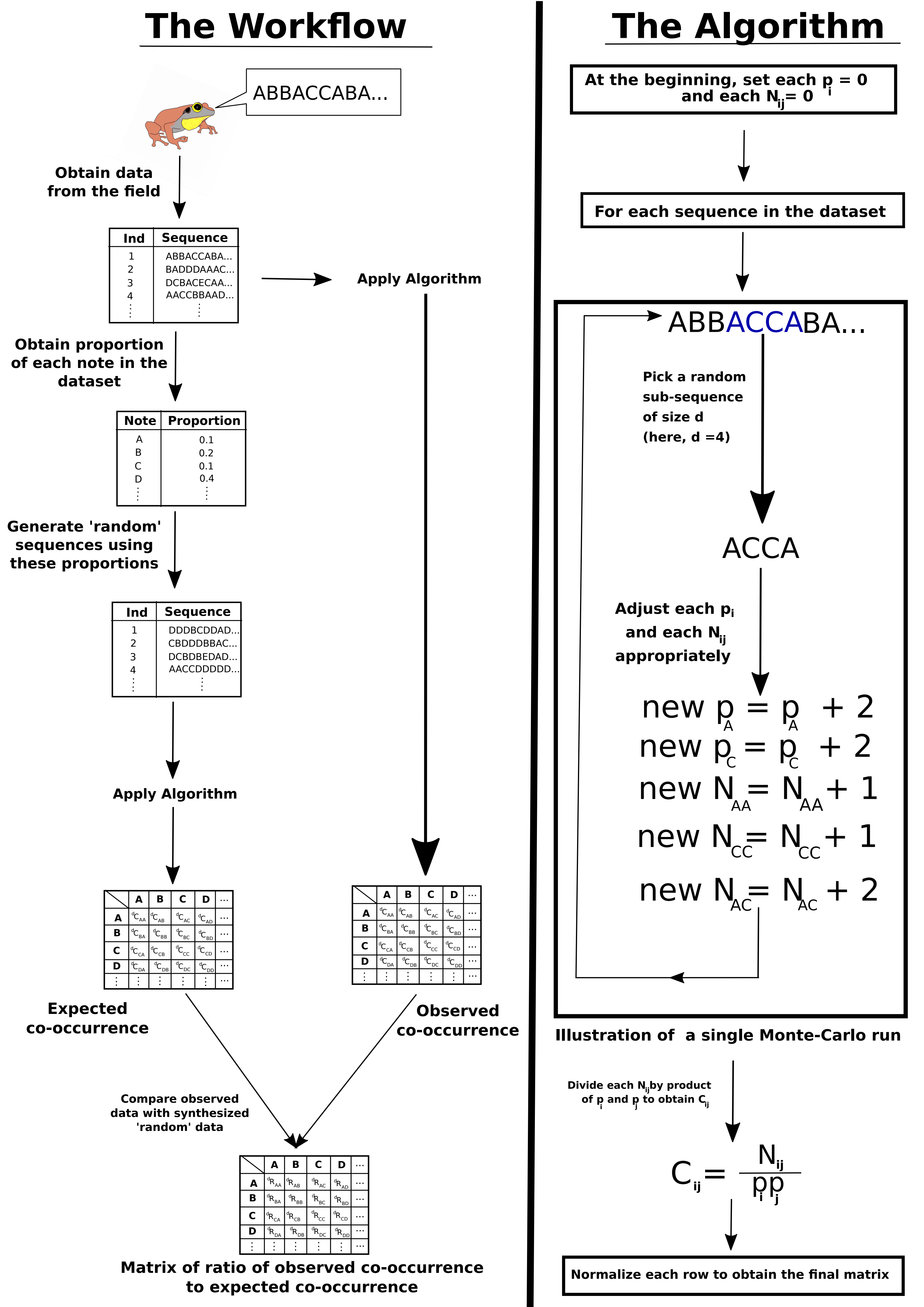

### Supplementary File 01

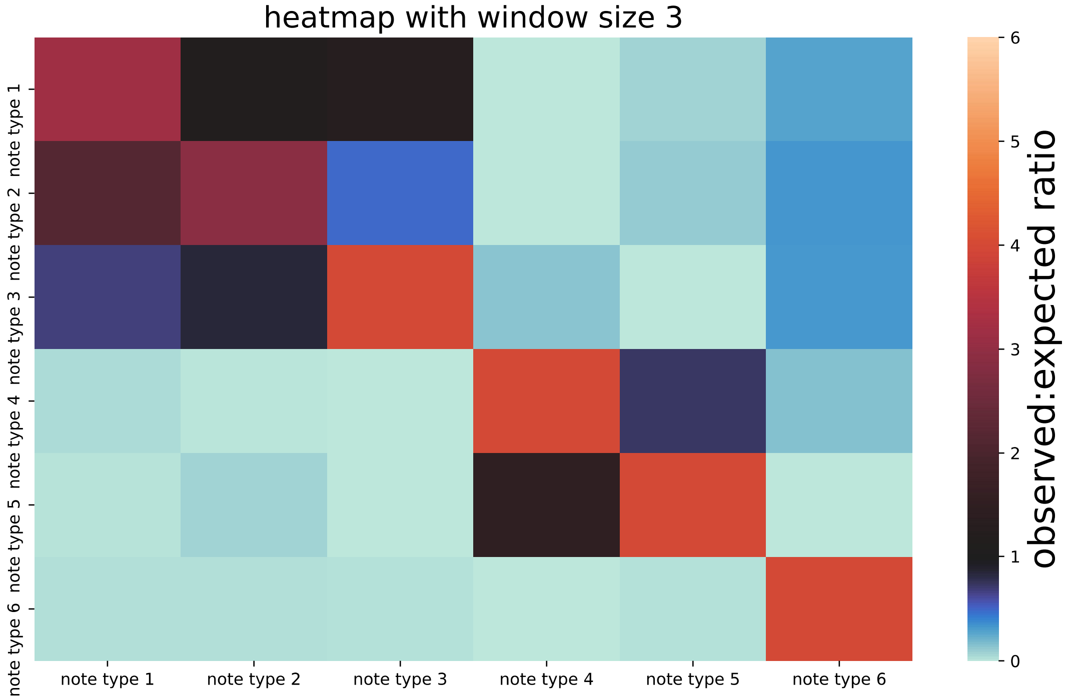

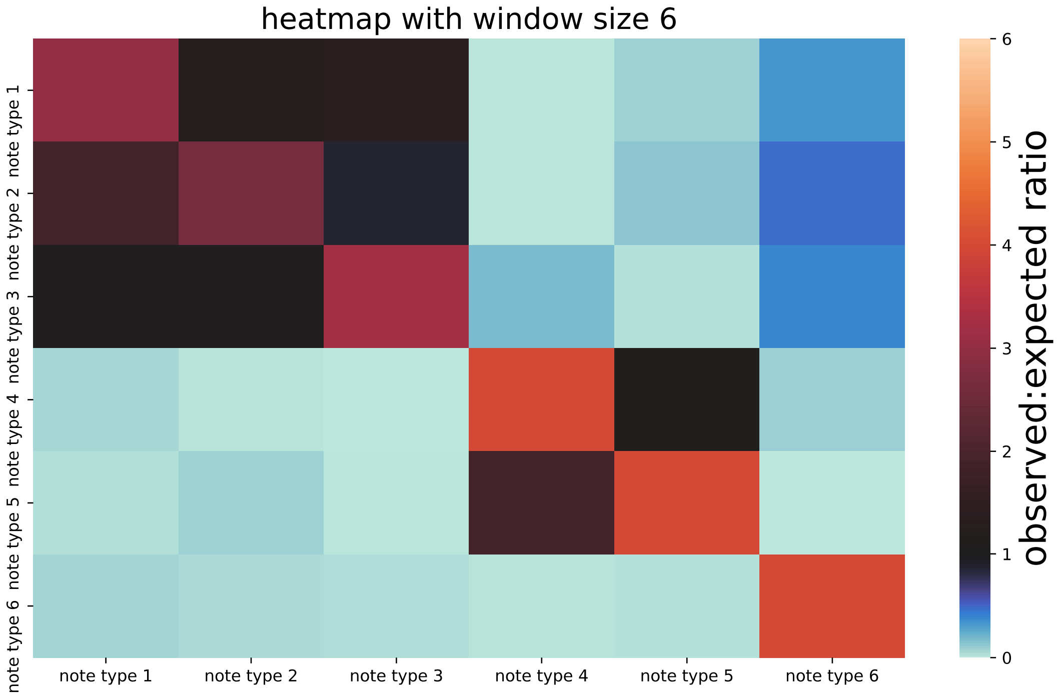

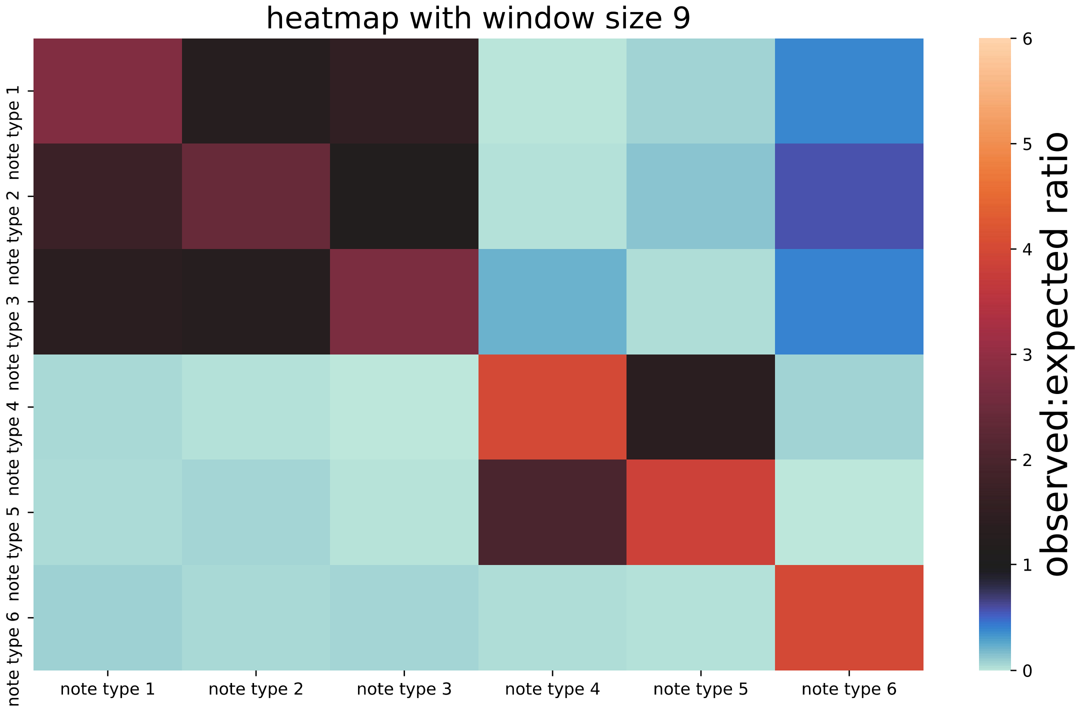

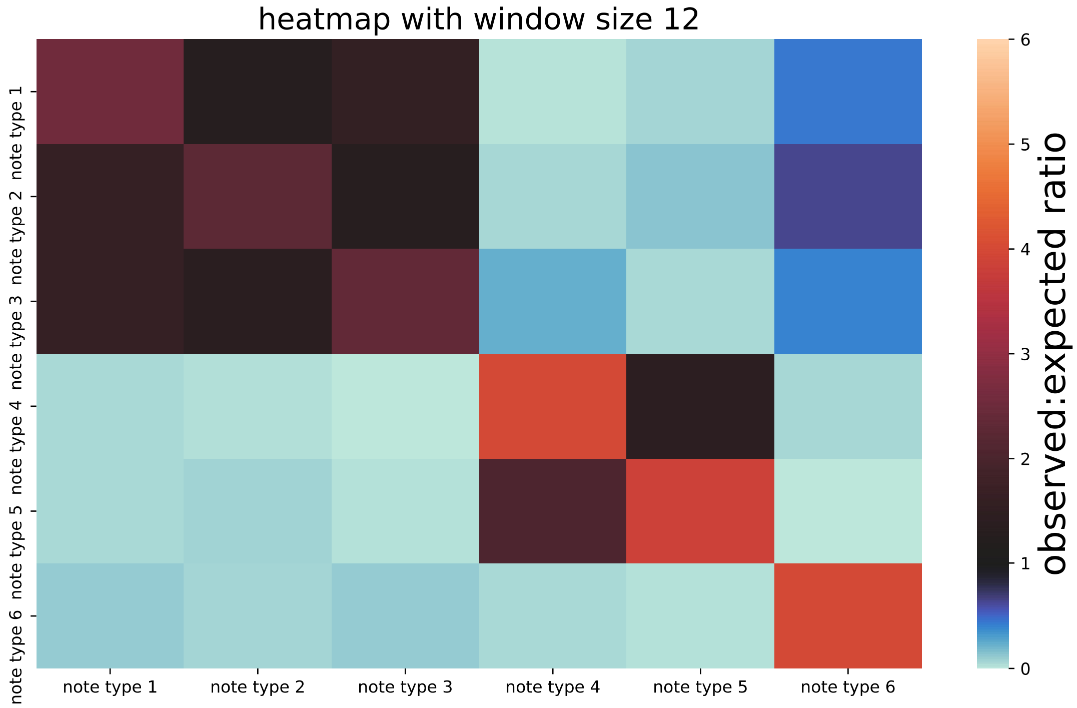
